# Seamless Assembly of Biological *<u>Parts</u>* into Functional *<u>Devices</u>* and Higher Order *Multi-Device <u>Systems</u>*

**DOI:** 10.1101/346080

**Authors:** Jeffrey C Braman, Peter J Sheffield

## Abstract

A new method is described for the seamless assembly of independent, prefabricated and functionally tested blunt-end, double strand nucleic acid *<u>parts</u>* (DNA fragments) into more complex biological *<u>devices</u>* (vectors) and higher order <u>multi-device *systems*.</u> Individual *parts* include bacterial selection markers, bacterial origins of replication, promoters from a variety of different species, transcription terminators, shuttle sequences and a variety of “N” and “C” terminal solubility/affinity expression tags. Pre-assembly modification of *parts* with DNA modifying enzymes is not required. Seamless assembly of multiple parts is accomplished in a single step using a specialized thermostable enzyme blend in about 30 minutes. Combinatorial assembly of *parts* is an inherent feature of the new process, substantially simplifying *device* and *system* optimization. To underscore the utility of the new process, *parts* were assembled into several protein expression *devices* in order to identify the optimal expression construct for a model target gene, as an example of the utility of the assembly process, and a higher order multi-*device system* is also described, for the over-expression of a four-enzyme bio-synthetic pathway, and optimized for end-product accumulation in *E. coli* as a paradigm for how this assembly process could be used to address the assembly of more complex biological pathways.

## Introduction

The discipline of synthetic biology has greatly benefitted from key enabling technologies such as DNA synthesis and sequencing becoming accessible to more researchers due to the reduction in the previously prohibitive financial entry point. To date however, a third enabling technology, molecular cloning, has not kept pace with technological advances made in DNA synthesis and DNA sequencing. One of the most highly recognized collection of techniques and materials developed to improve conventional cloning of biological *parts*, *devices* and *systems* is “BioBricks” (1, 2). Briefly, the “bricks”, or *parts,* of this technology represent cloned DNA sequences possessing defined functions, such as antibiotic resistance and ribosome binding sites. *Parts* are assembled to create larger *devices* such as protein expression vectors and several *devices* are joined into a *system* such as a biosynthetic pathway. BioBrick *devices* and *systems* are constructed by “hierarchical binary assembly” of *parts,* or “one-brick-at-a-time.” More specifically, BioBrick’s represent functional double strand DNA molecules housed within carrier plasmids flanked by universal and precisely defined upstream and downstream sequences that are technically not part of the BioBrick. These universal sequences contain restriction enzyme recognition sites for one of two closely related enzymes, each having slightly different recognition sequences but upon cleavage generate identical termini (isocaudomers). Linking two BioBricks together requires isolation of the individual *parts* from their carrier plasmids by specific isocaudomer(s) digestion, end repair in some cases, ligation and finally bacterial transformation. A major drawback to this technique is that BioBrick *parts* must not contain these restriction enzyme recognition sites within the sequences to be assembled. Also, BioBrick hierarchical binary assembly is time consuming, tedious and not conducive to combinatorial assembly.

Other assembly methods that convert *parts* into *devices* also rely on the isolation of *parts* and *devices* from dedicated BioBrick-like “destination vectors” (BioBricks [2], SLIC [3], Gibson [4], CPEC [5], SLiCE [6], and In-Fusion [http://www.clontech.com/US/Products/Cloning_and_Competent_Cells/Cloning_Resources/Selec-tion_Guides/In-Fusion_Cloning_Kits]). In other methods, significant *parts* manipulation with either one or more Type-II restriction enzymes is required (GoldenGate [7], MoClo [8], GoldenBraid [9]). Alternatively, *parts* manipulation with T5-exonuclease or a combination of Pfu and Taq DNA polymerases are required for Gibson (4) and DATEL (10) assembly methods, respectively, to create overlaps for subsequent annealing and ligation. In summary, assembly methods are complicated when restriction enzyme specificity must be considered at each stage of *parts* and *devices* design. Also, creating small *parts* between 50 and 250 base pairs with one or more enzymes possessing exonuclease activity is difficult due to the propensity of these enzymes to completely degrade the *parts*. It is apparent that these limitations curtail combinatorial experimental design and significantly slow the process of identifying optimal *devices* and *systems*.

Providing prefabricated and functionally validated *parts* to researchers without the need for retrieval from destination vectors, combined with a seamless protocol conducive to combinatorial assembly of *parts* into *devices* and higher order *systems,* would represent a significant improvement in synthetic biology molecular cloning. This paper describes such a system. Prefabricated *parts* are provided such that a wide variety of *devices* can be rapidly assembled. Appropriately chosen *parts* are combined and assembled in a single-tube reaction creating molecules that transform/transfect and properly function as *devices* in *E. coli*, mammalian and yeast (*S. cerevisiae*) cells. Multiple assembles can be performed in parallel to generate a collection of unique *devices* that can be used in combination to optimize novel biological *systems*. The utility of this technology, referred to as “SureVector” (SV), was validated by assembling *parts* into a collection of *devices* (plasmids) in order to perform a protein expression screen to identify the best expression tag combination for a model gene of interest (GOI). To further demonstrate the applicability of the SV process, a multi-*device system* was designed and constructed to reconstitute a four-enzyme biosynthetic pathway in *E. coli*. This multi-*device system* enabled the identification of several over-expressing bacterial clones within one week.

## Materials and Methods

### SureVector (SV) P*arts*

SureVector *parts* include bacterial origins of replication [OR], bacterial selectable markers [SM], XP-1 and XP-2 referred to as “Expansion *parts*“ containing sequences allowing replication and selection in a variety of organisms (*Saccharomyces cerevisiae* and mammalian cells; defined in more detail below), promoters [P] (*E. coli*, *Saccharomyces cerevisiae* and mammalian), and protein expression tags [T] (both C-terminal and N-terminal tags are available). Each *part* is flanked by unique 30 base pair (bp) sequences not found in any global DNA databases, such as NCBI’s Basic Local Alignment Search Tool BLAST <u>(http://blast.ncbi.nlm.nih.gov/Blast.cgi),</u> and allow specific assembly of *parts* into *devices* (example shown in Table 1) and ultimately *systems*.

**Table 1.**
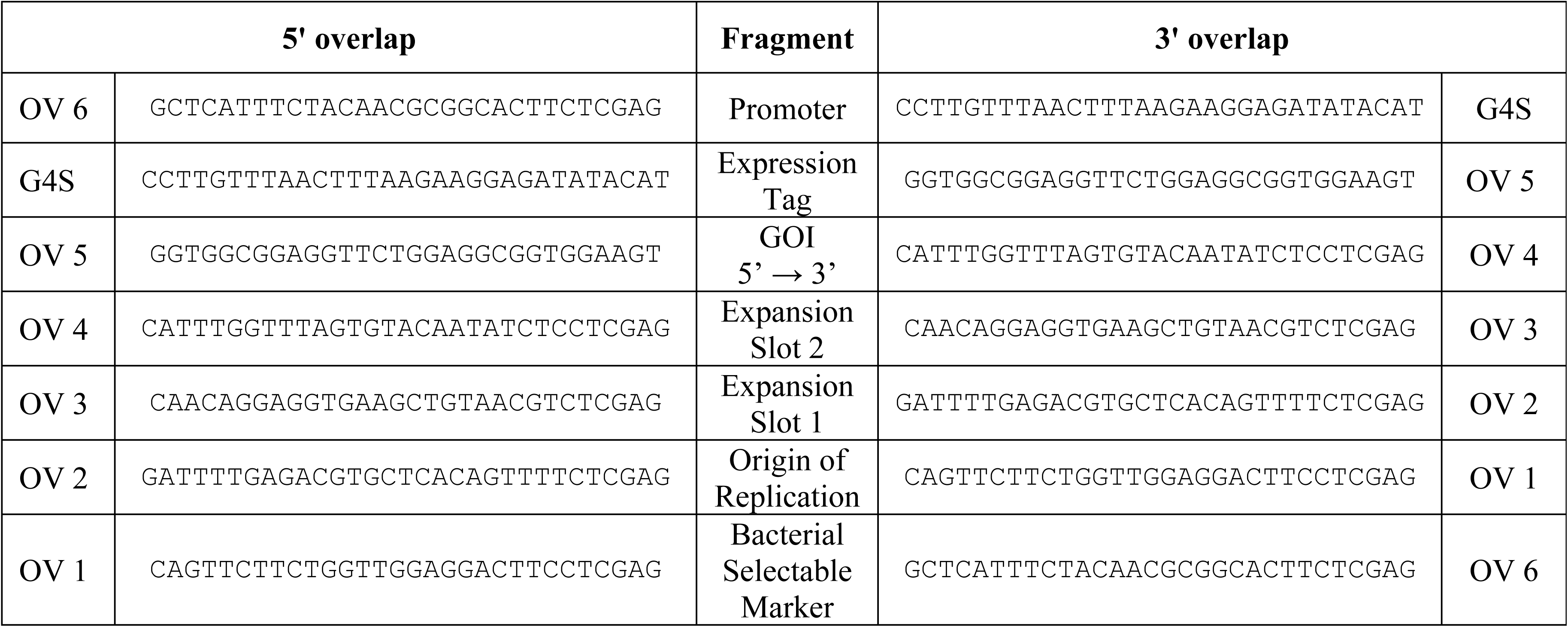
SureVector Overlaps (N-terminal Fusions)

### XP1 and XP2 Expansion *Parts*

XP1 *parts* contain either the yeast autonomous replication sequence (yARS or 2-micron circle) allowing plasmid replication in *S. cerevisiae* or a linker derived from a unique nucleotide sequence not possessing a function other than to tether a chosen bacterial origin of replication to an XP2 fragments during *parts* assembly.

The XP2 *parts* include either a non-functional tether sequence, as above, or the lacI repressor found in *E. coli*, or mammalian selection markers or yeast auxotrophic markers, allowing yeast to grow in the absence of an essential amino acid. XP2 *parts* designed specifically for use in mammalian cells include blasticidin, puromycin or hygromycin selection markers. For yeast, XP2 *parts* include the auxotrophic markers URA3 and HIS3 and the selection marker for hygromycin resistance. XP2 *parts* also contain a transcription terminator; either the bovine growth hormone polyA signal (bGH pA) specific for mammalian cells or a rho independent terminator for bacterial transcripts provided by the strong hairpin forming sequence (5’- GCCGCCAGCGGAACTGGCGGC-3’). These terminator sequences are positioned and oriented within the XP2 *part* to terminate transcripts from an upstream GOI.

### Large Scale Production and Purification of SV *Parts*

Large scale *parts* synthesis was performed by PCR in 96-well plates using sequence verified master plasmid templates. Each reaction contained 1 ng of plasmid template, 1x Herculase II reaction buffer (Agilent Technologies, Inc.), 0.25 mM of each dNTP, 0.4 µM of each PCR primer and 2 µl of pre-formulated Herculase II enzyme (Agilent Technologies, Inc.) in a final volume of 100 µL. Thermocycling conditions were: 1 cycle at 95 °C for 2 min.; 30 cycles at 95 °C for 20 sec., 55 °C for 20 sec. and 72 °C for 30 sec.; 1 cycle at 72 °C for 3 min. Contents of multiple 96-well plates for each SV *part* were pooled and purified using AMPure XP magnetic beads according to the manufacturer’s instructions (Beckman-Coulter). Correct lengths and purities of SV *parts* were assessed with an Agilent Technologies, Inc. BioAnalyzer.

### General Assembly of SV *Parts* into *Devices*

SV *parts* designed to assemble into a desired *device* were combined with a SV adapted GOI part and other reaction components were added as follows: 1x SureVector reaction buffer, 0.25 mM of each dNTP, 5.0 nM of each *part* (e.g. SM + OR + XP1 + XP2 + T + GOI + P) and 1 µL of pre-formulated enzyme in a final volume of 20 µL. Thermocycling of these components consisted of 1 cycle at 95 °C for 2 min.; 8 cycles at 95 °C for 20 sec., 55 °C for 20 sec. and 68 °C for 30 sec.; 1 cycle at 68 °C for 3 min. Following thermocycling, one unit of *Dpn* I restriction enzyme was added to the reaction and incubated for 5 minutes at 37 °C. One µL of the reaction was transformed into XL1-Blue Supercompetent *E. coli* cells (Agilent Technologies, Inc.) according to instructions and varying amounts (10, 20 and 50 µL) of the transformation mixtures were spread onto LB agar plates containing the appropriate antibiotic and incubated at 37 °C until colonies were easily visualized (12 – 16 hrs.). *Device* DNA was purified from select C-terminal reverse primer: colonies and either analyzed by restriction digestion, or sequenced to verify correct *parts* assembly, or used directly in downstream processes.

### SV *Parts* Assembly into Nedd5 Protein Expression *Devices*

To demonstrate an obvious use of the new SV cloning method, the human Nedd5 gene was chosen as a model GOI for performing an expression screening experiment. Nedd5 is a mammalian septin known to associate with actin-based structures such as the contractile ring and stress fibers and is involved in the process of cytokinesis in human brain tumors (11), although the specific nature of Nedd5 is not pertinent to this paper. The Nedd5 gene containing a start and a stop codon was adapted by PCR to be SV compatible by using the primers listed below:

N-terminal forward primer 5’- <u>GGTGGCGGAGGTTCTGGAGGCGGTGGAAGT***A***</u> ***TG***GGATCCATGTCTAAGCAACAACCAACTC-3’ and N-terminal reverse primer 5’-<u>TCGAGGAGATATTGTACACTAAACCAAATG</u>***TCA***CACATGCTGCCCGAGAGCCCCGCTGTCAC-3’.

The Nedd5 gene containing a start codon but lacking a stop codon was adapted by PCR to be SV compatible by using the following primers:

C-terminal forward primer:

5’- <u>CCTTGTTTAAACTTTAAGAGGAGGGCCACC</u>***ATG***GGATCCATGTCTAAGCAACAACCAACTC-3’ and

C-terminal forward primer:

5’-<u>CCACCGCCTCCAGAACCTCCGCCACCCACA</u>TGCTGCCCGAGAGCCCCGCTGTCACTGTCAC-3’.

Primers show the start codon or stop codon in bold-italicized type and all primers include unique 30 bases [underlined] for assembly with adjacent *parts*. The resulting Nedd5 PCR products were used as the model GOI and assembled into a variety of twelve expression constructs each containing a different C- or N-terminal expression tag. The following *parts* were used in these assemblies: Amp [SM] + pBR322 [OR] + XP-1 linker + XP-2 *lacI* + Nedd5 [GOI] with either C-terminal Tags[T] (c-Myc, thioredoxin, streptavidin binding protein, calmodulin binding protein, His6 or hemagglutinin) or N-terminal tags (GST, MBP, His6, SBP, CBP, or HisDbsA, calmodulin binding protein or hemagglutinin) + pTac [P] (Refer to section entitled “General assembly of SV *parts* into *devices*“).

### SV Nedd5 Expression *Devices* Screening

Nedd5 clones with different expression tags were cultured overnight at 37°C with shaking at 250 rpm in 1 ml of LB broth containing ampicillin (50 µg/ml). The following day, 10 ml cultures of LB broth containing ampicillin (50 µg/ml) were inoculated with these clones and incubated at 37°C with shaking at 250 rpm until the OD_600_, a measure of bacterial growth, reached 0.6 (approximately 1 hour). Protein expression from the P_Tac_ promoter was induced by the addition of IPTG to a final concentration of 0.5 mM followed by incubation for 20 hours at 30°C with shaking at 250 rpm. Volumes of cells equal to OD_600_ of 3.0 were removed from each culture at time zero (uninduced samples) and after 20 hours of incubation (induced samples). The samples were then centrifuged and cell pellets resuspended in 120 µl of 8M urea. The mixtures were vortexed well and incubated at 75°C for 5 min. Cell lysates were centrifuged and supernatants analyzed for Nedd5 expression by SDS-gel electrophoresis.

### SV Assembly of *Parts* and *Devices* into Higher Order *Systems e*xpressing DMRL

SV *parts* and *devices* were assembled to reconstitute the four enzyme *E. coli* biosynthetic pathway (*system*) for 6,7-<u>d</u>i<u>m</u>ethy-8-<u>r</u>ibityl<u>l</u>umazine (DMRL), the fluorescent precursor to riboflavin (rib pathway genes). The DMRL *system* construction was accomplished by first assembling *devices* with zero, one or two rib genes. PCR primers used to amplify rib open reading frame gene *parts* A (ribA – 591 bp), B (ribB – 654 bp), D (ribD – 1104 bp) and E (ribE - 471 bp) with appended SV overlaps to make them SV compatible were:

*E. coli* ribA – Gene ID = 945763

1. ribA_Forward Primer-N-Tag (56 bp):

ggtggcggaggttctggaggcggtggaagt***ATG***CAGCTTAAACGTGTGGCAGAAGC

2. ribA_Reverse Primer-RBS (67 bp):

gaaattgttaaattatttctagattcgaaaggagctcgaattcTTATTTGTTCAGCAAATGGCCCAT

*E. coli* ribB – Gene ID = 947526

1. ribB_Forward Primer-N-Tag (56 bp)

ggtggcggaggttctggaggcggtggaagt***A TG***AATCAGACGCTACTTTCCTCTTT

2. ribB_Reverse Primer-RBS (68 bp):

gaaattgttaaattatttctagattcgaaaggagctcgaattcTCAGCTGGCTTTACGCTCATGTGCC

*E. coli* ribD – Gene ID = 945620

1. ribD_Forward Primer-RBS (73 bp):

ttcgaatctagaaataatttaacaatttcacataaaggaggtaaata***ATG***CAGGACGAGTATTACATGGCGCG

2. ribD_Reverse Primer (54 bp)

ctcgaggagatattgtacactaaaccaaatgTCATGCACCCACTAAATGCAGGC

*E. coli* ribE - Gene ID = 946453

1. ribE_Forward Primer-RBS (74 bp):

ttcgaatctagaaataatttaacaatttcacataaaggaggtaaata***ATG***AACATTATTGAAGCTAACGTTGC

2. ribE_Reverse Primer: (55 bp)

ctcgaggagatattgtacactaaaccaaatgTCAGGCCTTGATGGCTTTCAATAC

Two sets of bi-cistronic *devices* were designed and assembled, one containing the ribA and ribD genes and the other containing the ribB and ribE genes. A ribosome binding site (RBS) was included in the 3’ region of the ribA and ribB genes downstream of their native stop codon and the same sequence was also included in the 5’ region upstream of the ATG start codon of the ribD and ribE genes. This RBS sequence was used as the overlap by which the ribD and ribE genes were positioned downstream of ribA and ribB genes, respectively. The intended outcome was to place two rib genes under control of one promoter and couple expression of the upstream and downstream rib genes via a second RBS between the two rib genes. This was done to attempt to balance expression levels of the individual rib genes. This second RBS overlap region between the rib genes was designed in such a way that the downstream rib gene was not in the same reading frame as the upstream rib gene thus preventing two gene products in the same *device* from becoming physically linked. An additional stop codon was also added to each upstream rib gene to further guard against translation read-through. Bi-cistronic vectors of this type have been used previously for preparation of nuclear receptor partners RAR and RXR (12) and for the analysis of NFΦB p50/p65 heterodimer (13). *Devices* lacking either the upstream or downstream rib gene *parts* were correctly assembled into circular molecules using 90 bp N-terminal and C-terminal “Non-Coding” linker *parts* NC-N and NC-C, respectively, and were made by overlap extension (14):

N-terminal “NC” replaces ribA or ribB *parts*.

rib - “NC” N-term_Forward ggtggcggaggttctggaggcggtggaagtgaaactgcactcatcgtccctcgaggagct

rib - “NC” N-term_Reverse gaaattgttaaattatttctagattcgaagagctcctcgagggacgatgagtgcagtttc

C-terminal “NC” replaces ribD or ribE *parts*.

rib - “NC” C-term_Forward ttcgaatctagaaataatttaacaatttcacataaaggaggtatagacagcatacgagtc

rib - “NC” C-term_Reverse ctcgaggagatattgtacactaaaccaaatgactcgtatgctgtctatacctcctttatg

Bi-cistronic *devices* that only contain a single rib gene required either a NC-N or NC-C *part* in lieu of the corresponding rib gene *part*. Three standard *parts* were used in both sets of rib *devices*; the T7 promoter-HIS6, XP1 linker and XP2 lacI. These 3 standard *parts* were used in various combinations with either the ampicillin or kanamycin selectable markers and the pBR322 or p15a bacterial origins of replication. Therefore, all SV rib *devices* were assembled from just seven SV *parts.* A total of 18 *device* level plasmids were constructed. These system level *devices* were designated by letter-number codes. “K” *devices* consisted of the kanamycin resistance marker (kan), the p15a origin of replication and either zero, one or two rib genes. “A” *devices* consisted of the ampicillin resistance marker (amp), the pBR322 origin of replication and either zero, one or two rib genes (Table 2). Higher order *systems* were created using various combinations of these *device* level plasmids by the co-transformation of two *devices.* For example, *devices* K6 and A7 resulted in co-expression of ribA-ribD genes from the K6 *device* and ribB-ribE genes from the A7 *device.* Combinations of *devices* were transformed into Agilent BL21(Gold) DE3 *E. coli* and spread onto LB- agar plates containing 100 µg/ml each of kanamycin and ampicillin (LB-kan-amp) plus 0.5 mM IPTG. Plates were incubated at 37°C for 12 to 18 hours and examined under UV light to identify DMRL expressing clones (*systems*) as evidenced by fluorescent colonies surrounded by fluorescent halos.

**Table 2.** SV *Devices* Required to Re-Create the DMRL Biosynthetic Pathway. Each *device* contains zero, one or two rib genes, and either the kanamycin resistance marker and p15a origin of replication prefabs (labeled “K”) or the ampicillin resistance marker and pBR322 origin prefabs (labeled “A”). Single rib gene control plasmids (K1 – K4; A1-A4) require either N- or C- terminal “non-coding” *parts* (NC-N and NC-C, respectively) to assemble complete *devices*. Zero rib gene control *devices* (K5 and A5) contain both NC-N and NC-C *parts* in place of both rib genes to assemble into complete *devices*. Co-transformation of one “K” *device* (e.g. K6 through K9) with one “A” *device* (e.g. A6 through A9) results in a higher order *system* expressing either 0, 1, 2, 3 or 4 rib genes.

| Kanamycin/p15a Rib “K” *Devices* |  |  |  |  |  |  |
| --- | --- | --- | --- | --- | --- | --- |
| *Device ID* | SM | Origin | Promoter | GOI1 | GOI2 | Type |
| K1 | Kan | p15a | T7-His | RibA | NC-C | Control |
| K2 | Kan | p15a | T7-His | RibB | NC-C | Control |
| K3 | Kan | p15a | T7-His | NC-N | RibD | Control |
| K4 | Kan | p15a | T7-His | NC-N | RibE | Control |
| K5 | Kan | p15a | T7-His | NC-N | NC-C | Control |
| K6 | Kan | p15a | T7-His | RibA | RibD | Test Plasmids |
| K7 | Kan | p15a | T7-His | RibB | RibE | Test Plasmids |
| K8 | Kan | p15a | T7-His | RibA | RibE | Test Plasmids |
| K9 | Kan | p15a | T7-His | RibB | RibD | Test Plasmids |

| Ampicillin/pBR322 Rib “A” *Devices* |  |  |  |  |  |  |
| --- | --- | --- | --- | --- | --- | --- |
| *Device ID* | SM | Origin | Promoter | GOI1 | GOI2 | Type |
| A1 | Amp | pBR322 | T7-His | RibA | NC-C | Control |
| A2 | Amp | pBR322 | T7-His | RibB | NC-C | Control |
| A3 | Amp | pBR322 | T7-His | NC-N | RibD | Control |
| A4 | Amp | pBR322 | T7-His | NC-N | RibE | Control |
| A5 | Amp | pBR322 | T7-His | NC-N | NC-C | Control |
| A6 | Amp | pBR322 | T7-His | RibA | RibD | Test Plasmids |
| A7 | Amp | pBR322 | T7-His | RibB | RibE | Test Plasmids |
| A8 | Amp | pBR322 | T7-His | RibA | RibE | Test Plasmids |
| A9 | Amp | pBR322 | T7-His | RibB | RibD | Test Plasmids |

**Table 3.** Combinations of Table 2 Functional and Control *Devices* Used to Re-Create the DMRL Biosynthetic Pathway *System*.

| Transformations into BL21(GOLD) |  |  |  |  |  |
| --- | --- | --- | --- | --- | --- |
| *System* |  | rib Genes Expressed | Rib genes Missing | Plate on | Assay |
| *Device 1* | *Device 2* |  |  |  |  |
| K1 | - | A | B, D and E | Kan | Control |
| K2 | - | B | A, D and E | Kan | Control |
| K3 | - | D | A, B and E | Kan | Control |
| K4 | - | E | A, B and D | Kan | Control |
| K5 | - | - | A, B, D and E | Kan | Control |
| K6 | - | A and D | B and E | Kan | Control |
| K7 | - | B and E | A and D | Kan | Control |
| K8 | - | A and E | B and D | Kan | Control |
| K9 | - | B and D | A and E | Kan | Control |
| A1 | - | A | B, D and E | Amp | Control |
| A2 | - | B | A, D and E | Amp | Control |
| A3 | - | D | A, B and E | Amp | Control |
| A4 | - | E | A, B and D | Amp | Control |
| A6 | - | A and D | B and E | Amp | Control |
| A7 | - | B and E | A and D | Amp | Control |
| A8 | - | A and E | B and D | Amp | Control |
| A9 | - | B and D | A and E | Amp | Control |
| A5 | - | - | A, B, D and E | Amp | Control |
| K1 | A9 | A, B and D | E | Kan+Amp | Control |
| K2 | A8 | B, A and E | D | Kan+Amp | Control |
| K3 | A8 | D, A and E | B | Kan+Amp | Control |
| K4 | A9 | E, B and D | A | Kan+Amp | Control |
| K6 | A7 | A, D, B and E | - | Kan+Amp | Test |
| K7 | A6 | B, E, A and D | - | Kan+Amp | Test |
| K8 | A9 | A, E, B and D | - | Kan+Amp | Test |
| K9 | A8 | B, D, A and E | - | Kan+Amp | Test |

### Validation of DMRL *System* Synthesis - Assay, Purification and MS Analysis

Monitoring clones for DMRL synthesis on agar plates was straightforward as the DMRL fluoresces within colonies and is secreted into the surrounding media creating fluorescent halos around DMRL positive colonies. DMRL production from clones cultured in liquid media was also straightforward as the compound possesses a characteristic visible light absorption spectrum with λ max OD_490_ (15) that can be measured in cell free supernatants. Validation of DMRL synthesis required its production, purification and analytical characterization. Clone K6A7 produced pronounced fluorescent colony-halos and was chosen for this purpose. A single colony was inoculated into 3 ml of LB-kan-amp liquid media and incubated overnight at 37°C with shaking at 250 rpm. A 2.0 ml sample of this culture was inoculated into a 500 ml Erlenmeyer flask containing 100 ml of LB-kan-amp and incubation continued as before until the OD_600_ value reached 0.35. IPTG was added to a final concentration of 0.5 mM and incubation continued for an additional 18 hours. Cells were removed by centrifugation and acetic acid added to the supernatant to a final concentration of 5%. This sample was applied to a 2.5 × 3.5 cm column of Florisil (Sigma-Aldrich) equilibrated with 5% acetic acid. The column was washed with one liter of 5% acetic acid and DMRL eluted with 100 ml of 3% pyridine. Solvent was removed by evaporation and the residue suspended in 5 ml of water. This sample was applied to a 2.5 × 35 cm column of chromatography grade cellulose (Sigma-Aldrich) equilibrated with water and DMRL was eluted with water. Forty milligrams of DMRL were recovered from 0.8g of cells (wet weight) and was analyzed by mass spectrometry. The sample was run on an Agilent 6538 QTOF coupled to an Agilent 1100/1200 lc stack. The column was a Zorbax SB-C18 0.5 × 150 mm. Flowrate = 20µl/min. Injection volume = 5µl. Solvent A was H_2_O – 0.1% formic acid and solvent B was acetonitrile – 0.1% formic acid. Gradient: T0: 95% A - 5% B; T10: 75% A - 25% B; T12: 50% A - 50% B; T15: 5% A - 95% B; T20: off. Five min. re-equilibration time. For the ms run, a range from m/z = 85 to m/z = 1100 was scanned. For the ms/ms run, m/z 327.1 was targeted, with three different collision energies: ce = 10V, ce = 20V, and ce = 40V.

### Quantities and Rates of DMRL Synthesis from *Devices and Systems*

Single colonies from *devices* and *systems* listed in Table 2 were purified by re-streaking onto fresh LB-kan-amp plates without IPTG. Three colonies from each plate were cultured overnight at 37°C with shaking at 250 rpm in separate tubes containing 3 ml of LB-kan-amp. The next day, one hundred microliters of each culture were added to 4.9 ml of LB-kan-amp and incubated until OD_600_ reached between 0.3 and 0.5. IPTG was added to a final concentration 0.5 mM and incubation continued for three hours after which OD_600_ values were re-measured. Cultures were centrifuged to remove cells and OD_409_ values of supernatants obtained. OD_409_ values were normalized relative to OD_600_ values measured post IPTG addition and the resulting numbers compared. DMRL was synthesized exclusively by clones containing all four rib genes (two devices each expressing two rib genes) – *systems* K6A7, K7A6, K8A9, K9A8.

DMRL synthesis rates of *systems* K6A7, K7A6, K8A9, K9A8 and negative control K5A5 were obtained by inoculating single colonies into 5 ml of LB-kan-amp media followed by overnight incubation at 37°C with shaking at 250 rpm. One ml of these starter cultures was inoculated into separate 250 ml flasks each containing 49 ml of LB-kan-amp. Incubation at 37°C with shaking at 250 rpm continued until OD_600_ values reached between 0.3 and 0.5. IPTG was added to a final concentration of 0.5 mM and incubation continued. At regular intervals, 1.0 ml samples were retrieved from each culture and OD_600_ values measured. Samples were then centrifuged to remove cells and OD_409_ values of supernatants obtained. OD_409_ values were normalized relative to OD_600_ values measured post IPTG addition and the resulting numbers compared.

## Results

### Design Features of SV *Parts* and Assembly Process

The key design features of SV *parts,* ensuring precise and ordered joining, are the unique 30 bp sequences incorporated into each PCR primer used to generate a *part* (see Materials and Methods section). Figure 1 schematically represents how seven SV *parts* align and overlap due to this design feature.

**Figure 1.**
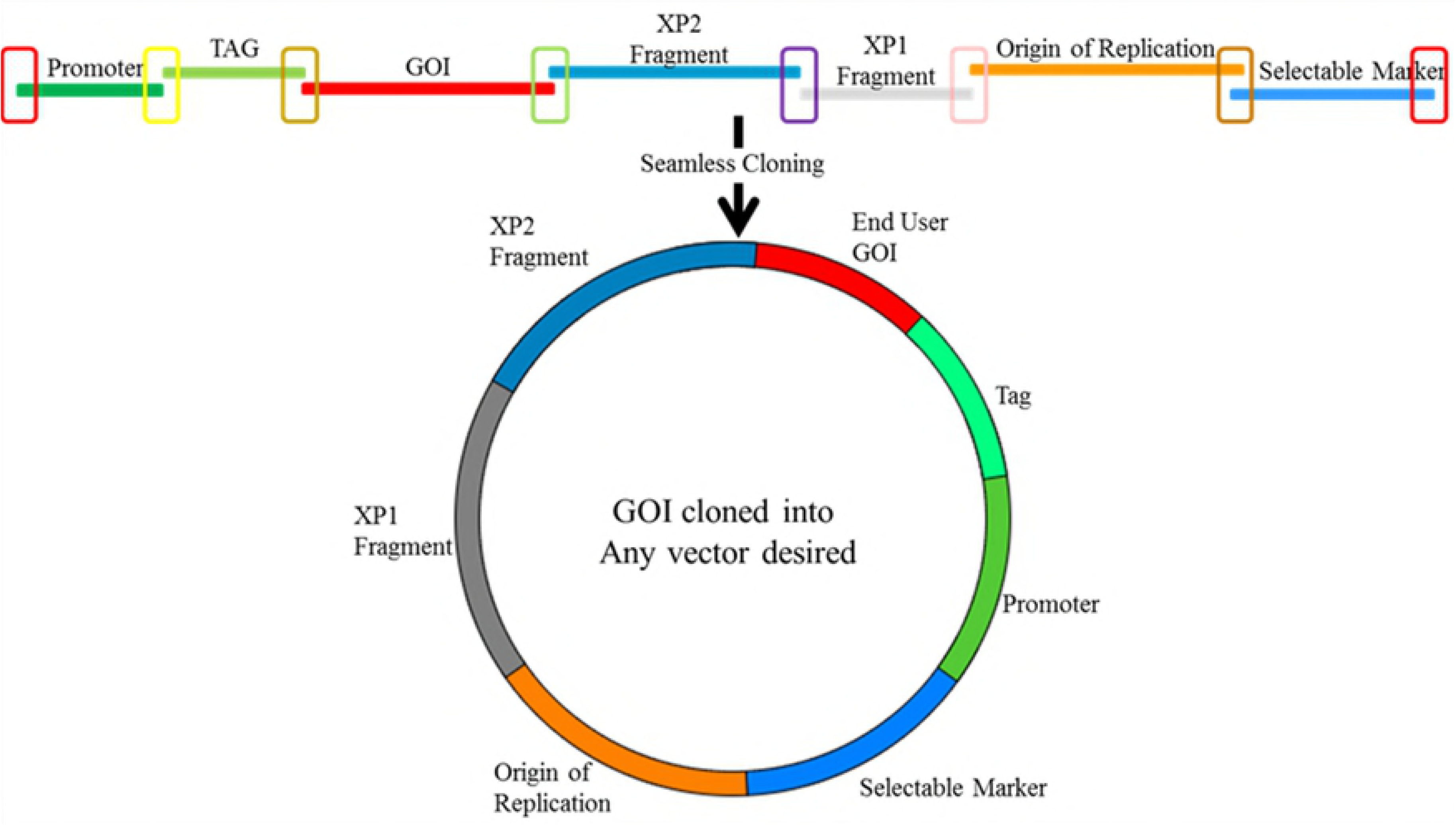
Schematic of a seven SV *part* assembly into a GOI Expressing *Device*. Different colored open rectangles highlight unique 30 bp overlaps between functional *parts.* Sequences represented by the two end rectangles (red) also overlap. The mixture of *parts* is treated with the SureVector enzyme assembly blend resulting in a *device,* represented by the closed circle that will transform, replicate and express a GOI in *E.coli.*

Assembling *parts* listed in

**Table 1** in the manner described in the Materials and Methods section and in the Figure 1 legend generated *devices* expressing GOI’s with an N-terminal fusion protein: -Bacterial Selectable Marker-Bacterial Origin of Replication-XP1-XP2-GOI←Expression TagFPromoter-(← denotes direction of promoter and gene expression)

The assembly mechanism of linking *parts* into higher order *devices* is represented in Figure 2. *Parts* are denatured and adjacent *parts* anneal due to the 30 bp overlaps. Exposed 3’-OH ends are partially extended by a polymerase resulting flaps that are digested by an endonuclease and covalently joined by a ligase.

**Figure 2.**
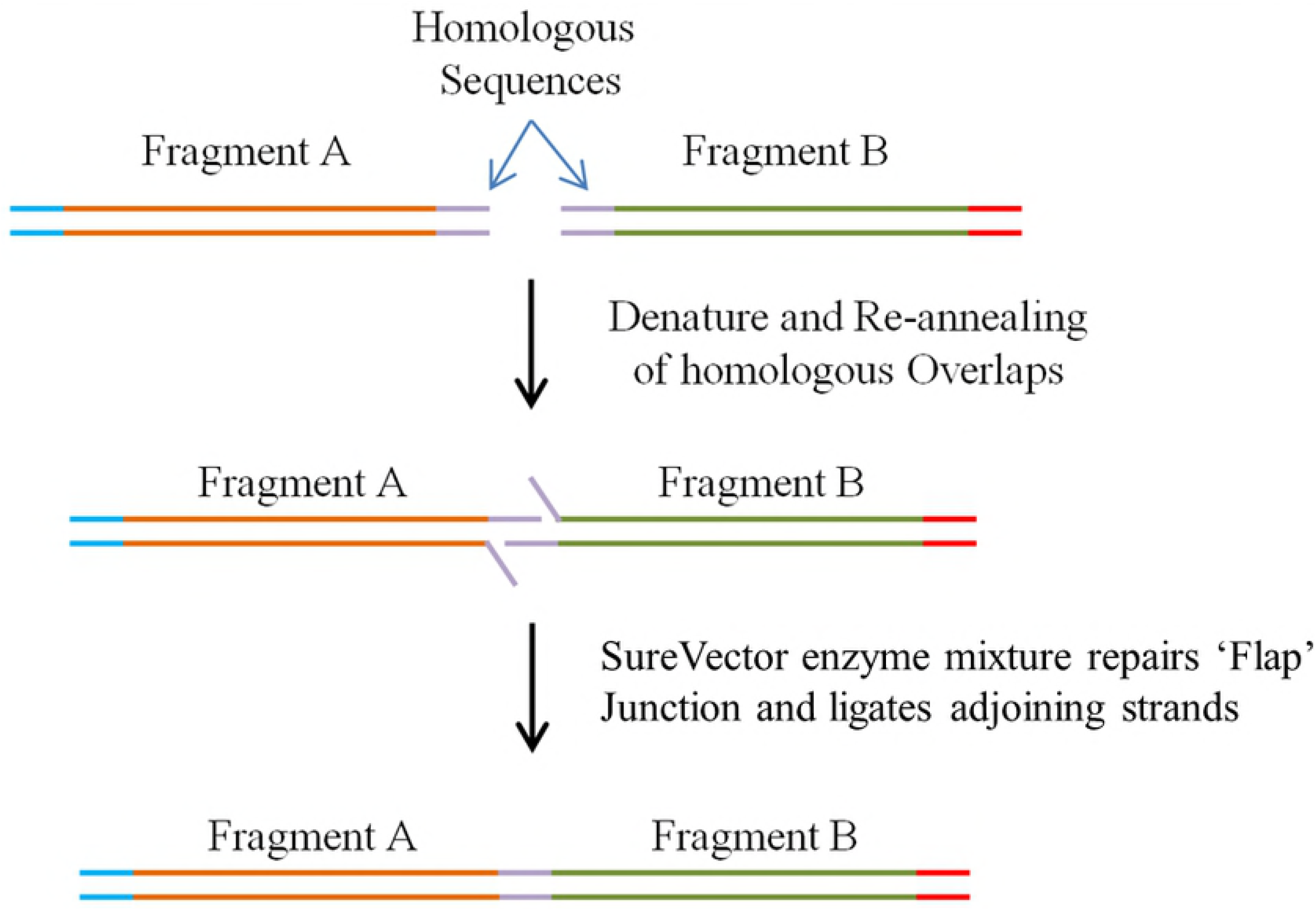
Schematic Showing How Adjacent SV P*arts* are Assembled; *Parts* A and B Possess Homologous Ends. Following denaturation and annealing, resulting free 3’ ends are extended, “flaps” digested and the two parts ligated.

Figure 3 represents the combinatorial assembly power of the SV process. Different functional *parts* are rapidly assembled into multiple configurations in parallel assembly experiments to determine the best organization for, in this case, expression of single GOI.

**Figure 3.**
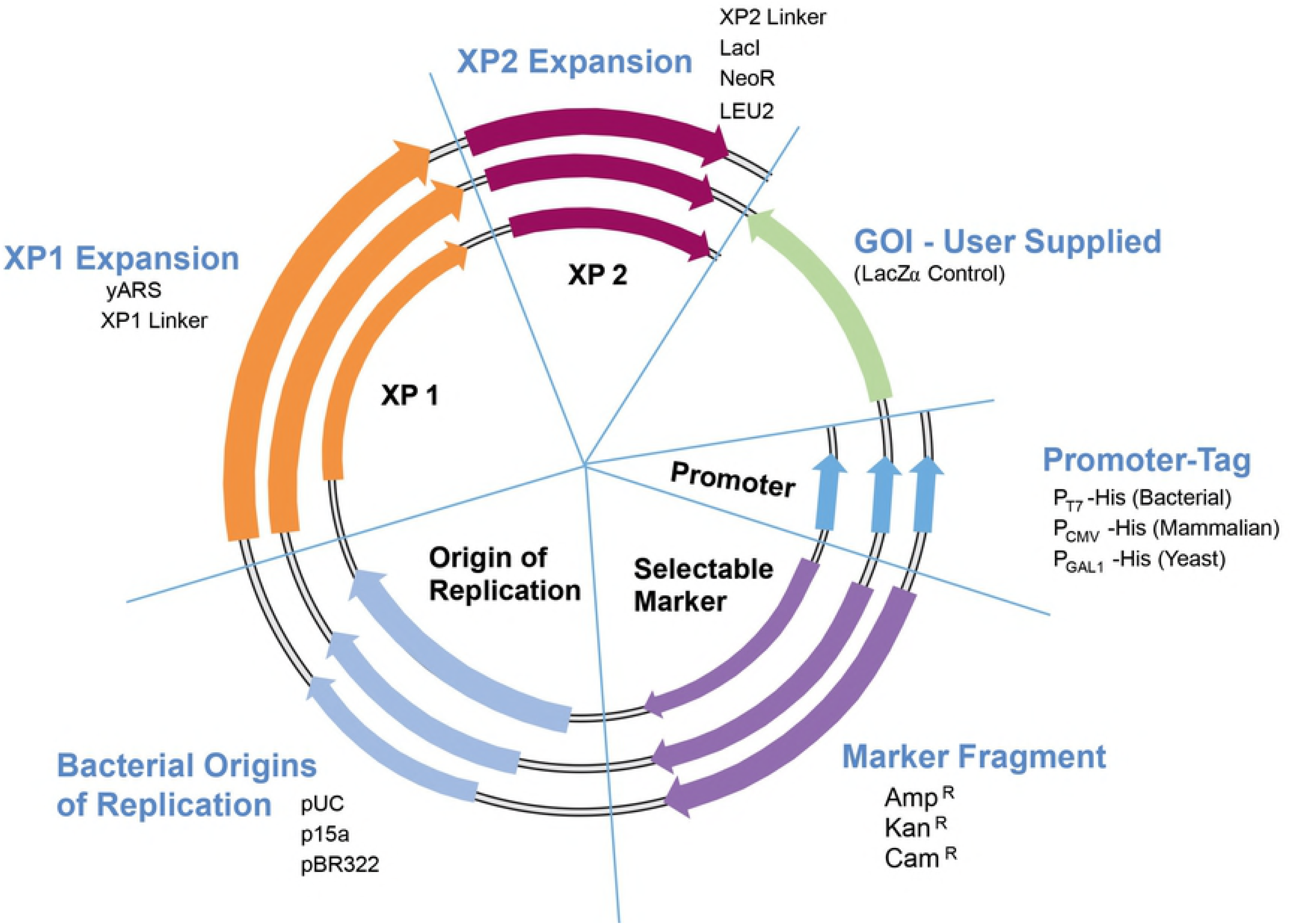
Collection of SV *Parts* and Assembly Design. A variety of *parts* and GOI’s can be assembled into many functional *devices.*

### SV Nedd5 Expression *Devices* Screening

SDS-page analysis of twelve Nedd5 expression *devices,* fused with N- or C- terminal tags identified the best expression constructs (green arrows shown in Figures 4a and b highlight proteins from induced cultures). Quantities of expressed fusion proteins varied with the type of expression tag and were highest with N-terminal tagged MBP/Nedd5, HisDbsA/Nedd5, and CBP/Nedd5 *devices* and C-terminal tagged Nedd5/SBP, Nedd5/c-myc and Nedd5/His6 *devices.* Lesser quantities of fused proteins were expressed as N-terminal tagged GST/Nedd5 and His6/Nedd5 *devices* and C-terminal tagged Nedd5/Thioredoxin and Nedd5/CBP *devices.* The least quantities of fusion Nedd5 were N-terminal tagged SBP/Nedd5 and C-terminal tagged Nedd5/HA *devices*. It is worth emphasizing that this protein expression screening experiment, starting from assembly of *parts* into *devices* and analyzing protein expression was completed in less than three days.

**Figure 4a.**
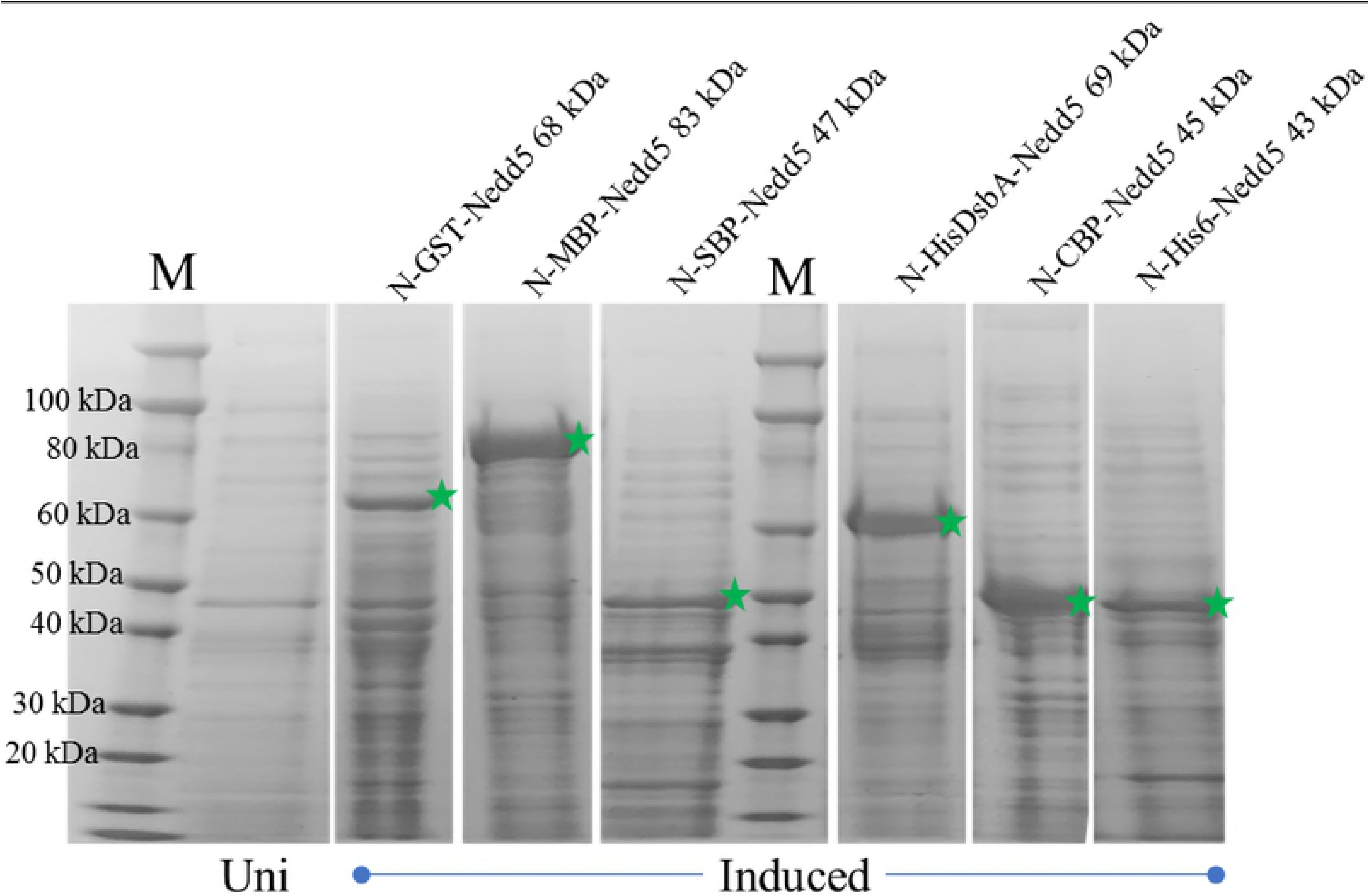
SDS-PAGE Gel of SV Nedd5 Expression *Devices* with Different N-Terminal Tags. M = protein molecular weight marker; Uni = Uninduced sample; Induced samples. Green stars (*) denote expressed N-terminal tagged Nedd5-fusion proteins.

**Figure 4b.**
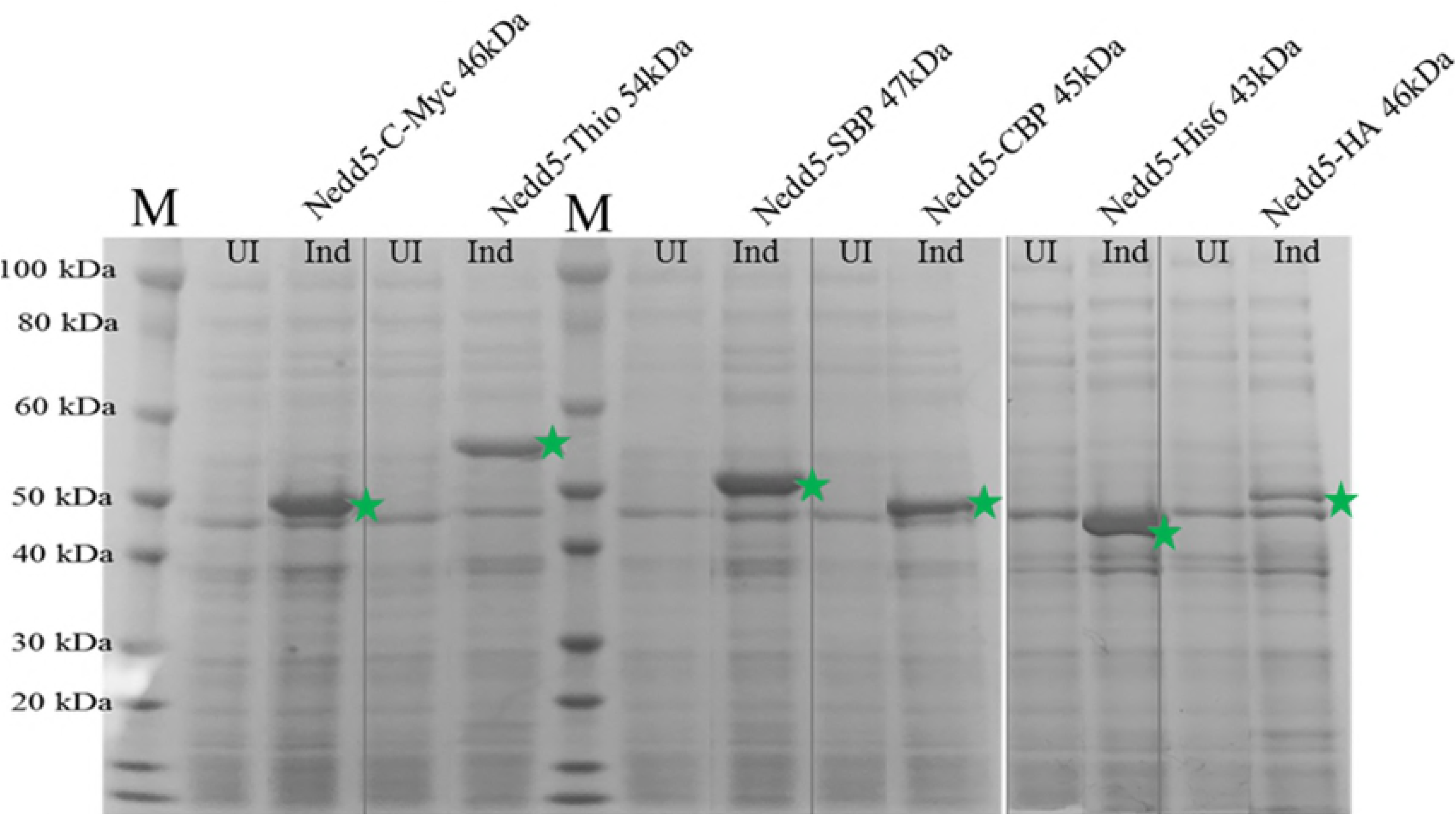
SDS-PAGE Gel of SV Nedd5 Expression *Devices* with Different C-Terminal Tags. M = protein molecular weight marker; UI = Uninduced samples; Ind = Induced samples. Green stars (*) denote expressed C-terminal tagged Nedd5-fusion proteins.

### Re-Creating a Biosynthetic Pathway

The utility of SV *parts* assembly into *devices* and ultimately higher order *systems* was demonstrated by recreating the *E. coli* biosynthetic pathway for 6,7-<u>d</u>i<u>m</u>ethy-8-ribityl<u>l</u>umazine (DMRL), the fluorescent precursor to riboflavin (Figure 5; 16). DMRL is synthesized by four unique enzymes (expressed from ribA, ribB, ridD and ribE genes) and substrates. The critical initial substrates for this pathway are GTP and ribulose-5’-phosphate (R5P). They are funneled through a “nucleotide conversion route” (NCR), a “sugar conversion route” (SCR) and a “converging condensation route” (CCR). The NCR starts by the hydrolytic removal of carbon atom 8 from the imidazole ring of GTP by GTP Cyclohydrolase II (ribA gene product; 17) yielding 2,5-diamino-6-ribosylamino-4(3H)-pyrimidinone-5’-phosphate (DARPP). DARPP is converted to 5-amino-6-ribitylamino-2,4(1H,3H)-pyrimidinedione-5’-phosphate (ARPP) by diaminohydroxyphosphoribosylaminopyrimidine deaminase/5-amino-6-ribitylamino-2,4(1H,3H)-pyrimidinedione reductase (ribD gene product; 18, 19), a “fused” enzyme possessing both NCR and SCR-relevant activities. As the enzyme name describes, the nucleotide base of DARPP is deaminated followed by reduction of the ribosyl moiety to ribityl with NADPH serving as reductant. Separately, L-3,4-dihydroxy-2-butanone-4-phosphate synthase, an enzyme exclusively in the SCR (ribB gene product; 20) converts ribulose-5’-phosphate (R-5-P) to L-3,4- dihydroxy-2-butanone-4-phosphate (DHBP) and formate. Both ARPP and DHBP are dephosphorylated and enter the CCR via DMRL synthase (ribE gene product; 21) resulting in DMRL.

**Figure 5.**
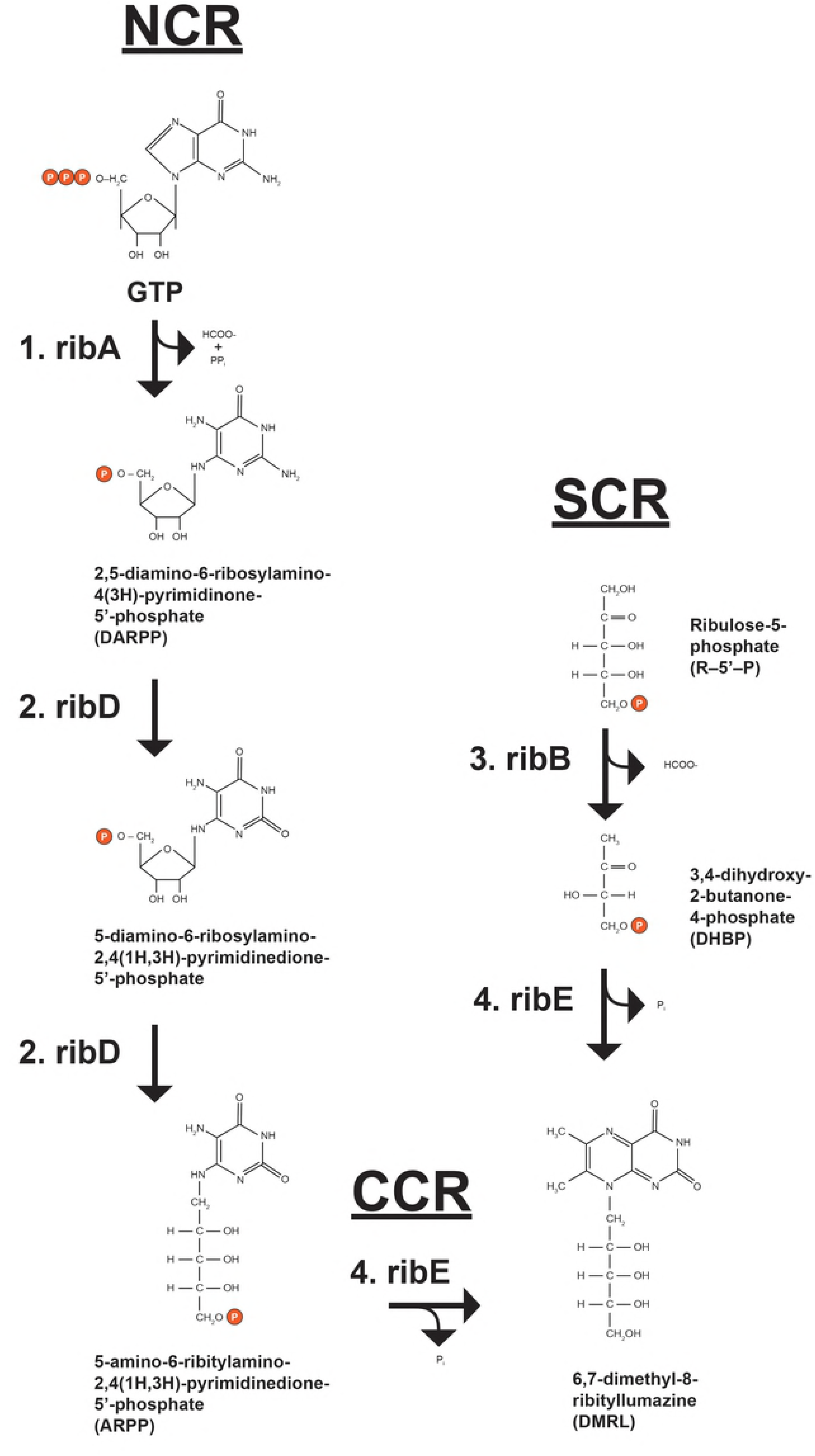
DMRL Biosynthetic Pathway. The imidazole ring of GTP is hydrolytically removed by GTP Cyclohydrolase II (rxn. 1; ribA) yielding 2,5-diamino-6-ribosylamino-4(3H)-pyrimidinone-5’-phosphate (DARPP), formate and pyrophosphate. DARPP is converted to 5- amino-6-ribitylamino-2,4(1H,3H)-pyrimidinedione-5’-phosphate (ARPP) by fused diaminohydroxyphosphoribosylaminopyrimidine deaminase/5-amino-6-ribitylamino2,4(1H,3H)-pyrimidinedione reductase (rxns. 2; both ribD). Separately, ribulose-5-phosphate (R5P) is converted to L-3,4-dihydroxy-2-butanone-4-phosphate (DHBP) and formate by L-3,4- dihydroxy-2-butanone-4-phosphate synthase (rxn. 3; ribB). ARPP and DHPB are dephosphorylated and then condensed by DMRL synthase (rxn. 4; ribE) producing 6,7-dimethyl8-ribityllumazine (DMRL).

### Pathway Construction

DMRL pathway construction was accomplished by first assembling SV *parts* into *devices* (Table 2) containing zero, one and two (“bi-cistronic”) rib genes. PCR primers used to amplify rib gene *parts* A, B, D and E are shown in Materials and Methods. They allowed amplification of rib open reading frames ribA – 591 bp, ribB – 654 bp, ribD – 1104 bp and ribE - 471 bp with appended correct overlaps to make them SV compatible. Two sets of bi-cistronic *devices*, one containing the ribA and ribD genes and the other containing the ribB and ribE genes were also designed and assembled (Table 2). Ribosome binding sites (RBS) were designed and incorporated into the 3’and 5’ flanking regions of the rib genes and subsequently used as unique overlaps between the genes such that ribD and ribE genes were positioned downstream of the ribA and ribB genes, respectively. The intended outcome of placing two rib genes under control of one T7 promoter and coupling expression of the upstream and downstream rib genes was balanced expression levels. The RBS placed in the 5’ regions of both ribD and ribE genes promoted downstream rib gene translation efficiency. The RBS overlap between the upstream and downstream rib genes was designed so that the downstream rib genes were out of frame with the upstream rib genes thus preventing two gene products in the same *device* from becoming physically linked. An additional stop codon was also added to each upstream rib gene as an added prevention to translation read-through. Bi-cistronic *devices* of this type have been used previously for preparation of nuclear receptor partners RAR and RXR (13) and for the analysis of NFΦB p50/p65 heterodimer (14). SV *devices* lacking one or both rib genes were also assembled using N-terminal and C-terminal “Non-Coding” *parts*; NC-N and NC-C, respectively (Table 2). Zero or single rib gene control *devices* required either an NC-N or NC-C *part*, or both in the case of the double negative control, in lieu of a rib gene *part*. Common SV *parts* used in both sets of rib *devices* were the T7 promoter-HIS_6_, XP1 linker and XP2 lacI (allowing IPTG induction of rib gene *devices* expression) in addition to various combinations of ampicillin and kanamycin selectable markers with pBR322 or p15a bacterial origins of replication. Therefore, all 18 SV rib *device* plasmids were assembled from just 7 seven SV *parts*, not including the rib gene *parts*.

Table 2 *devices* were transformed either individually or in various co-transformation scenarios (Table 3) into Agilent BL21(Gold) DE3 *E. coli* and spread onto LB- agar plates containing 100 µg/ml each of kanamycin and ampicillin (LB-kan-amp) plus 0.5 mM IPTG. Plates were incubated at 37°C for twelve to eighteen hours. All colony *devices* and *systems* were easily monitored for DMRL synthesis by irradiating plates with UV light and visualizing fluorescent colonies surrounded by fluorescent halos. DMRL producing colony *systems* contained all four rib genes (Figure 6 - left panel). Colonies containing only 1, 2 or 3 rib genes, as well as the double negative control lacking all 4 rib genes (clone K5A5 assembled with NC-N and NC-C *parts*), did not produce DMRL (Figure 6 - right panel; no fluorescent colonies and halos).

**Figure 6.**
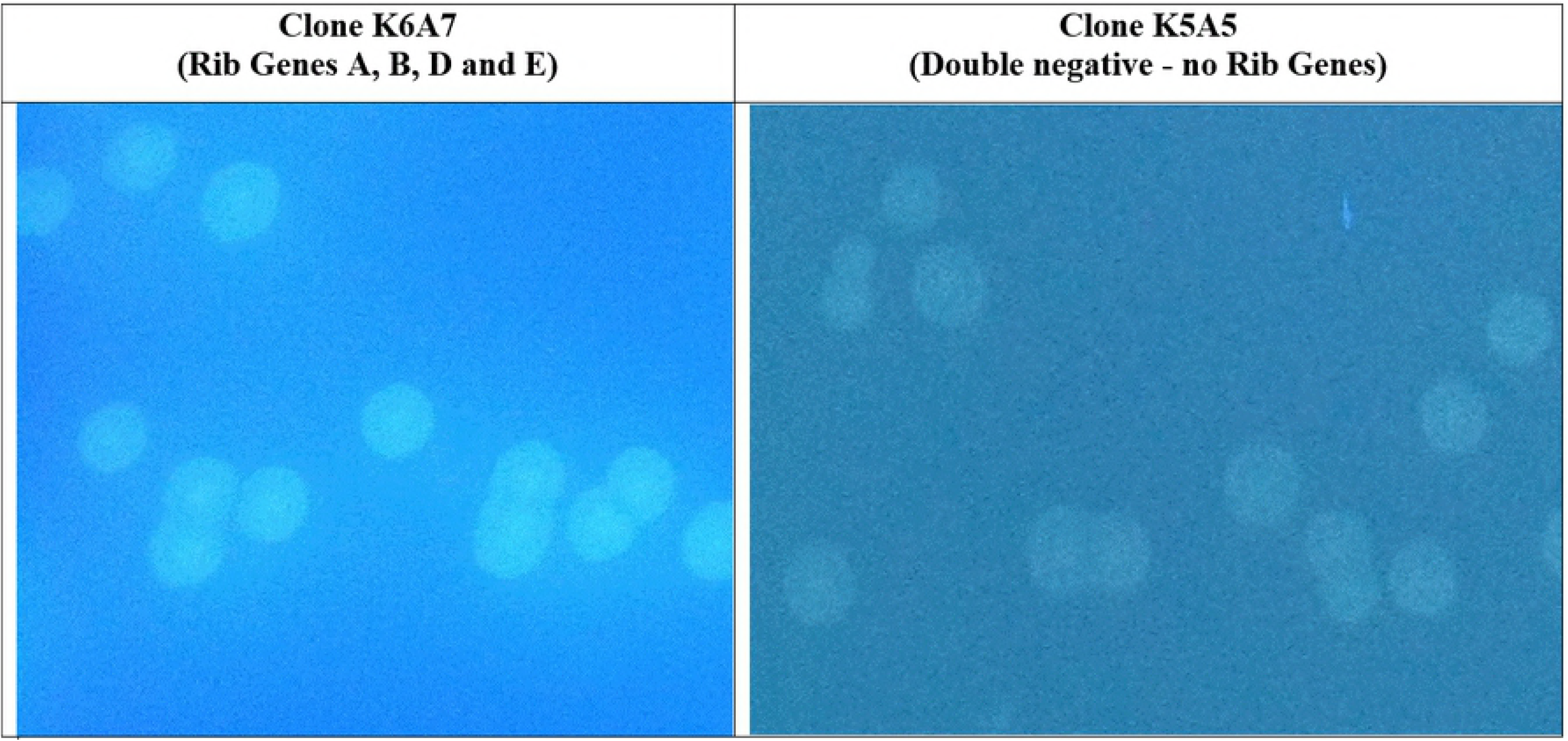
Co-Transformation of Two SV Compatible *Devices* Containing All 4 rib Biosynthetic Genes Results in DMRL Synthesis. *Devices* K6 and A7 (see Table 2) were co-transformed into Agilent BL21(Gold)DE3 *E.coli* and spread onto LB-kan-amp-IPTG plates. Resulting colonies were examined under unfiltered UV light. All resulting K6A7 *systems* contained four rib genes and produced DMRL as evidenced by fluorescent colonies with fluorescent halos. Control (*device* K5A5) through three rib genes (see Table 2) did not produce DMRL as no fluorescent halos were detected.

### Characterization of DMRL Synthesized by SV rib *System* K6A7

DMRL synthesized and purified from *system* K6A7 (Table 3) was characterized by mass spectrometry (see Materials and Methods). For the ms run, a range from m/z=85 to m/z=1100 was scanned. The theoretical m/z value for DMRL is 327.12990 and the observed m/z value was 327.12914 (data not shown). In addition, for the ms/ms run the m/z value of 327.1 was targeted, using three different collision energies at 10V, 20V, and 40V. The expected exact mass of fragmented DMRL was 192.06 and the observed value was 192.07 (data not shown) thus confirming that SV *system* K6A7 produced DMRL.

### Quantities and Rates of DMRL Synthesis from DMRL *Devices and Systems*

Figure 7a confirms that only SV *systems* K6A7, K7A6, K8A9, K9A8, each containing two rib gene *devices* expressing all four rib genes, produced DMRL. Clones K6A7 and K7A6 produced the highest levels of DMRL and most rapidly (Figures 7a and b). These *systems* are composed of d*evices* K6 and A6 expressing enzymes from ribA and ribD genes, respectively, both in the NCR path. ribA is positioned immediately downstream of the T7 promoter. *Devices* K7 and A7 express enzymes from ribB and ribE genes, respectively, with ribB in the SCR path and ribE in the CCR path. ribB is positioned immediately downstream of the T7 promoter. Total DMRL accumulation and slower synthesis rates are similar in *systems* K8A9 and K9A8. These *systems* are composed of d*evices* K8 and A8 expressing enzymes from ribA and ribE genes, respectively, with ribA in the NCR path and ribE in the CCR path. ribA is positioned immediately downstream of the T7 promoter. *Systems* composed of *devices* K9 and A9 express enzymes from ribB and ribD genes, respectively, with ribB in the SCR path and ribD in the NCR path. ribB is positioned immediately downstream of the T7 promoter. How these *system* configurations resulted in different totals and rates of DMRL synthesis were not examined.

**Figure 7.**
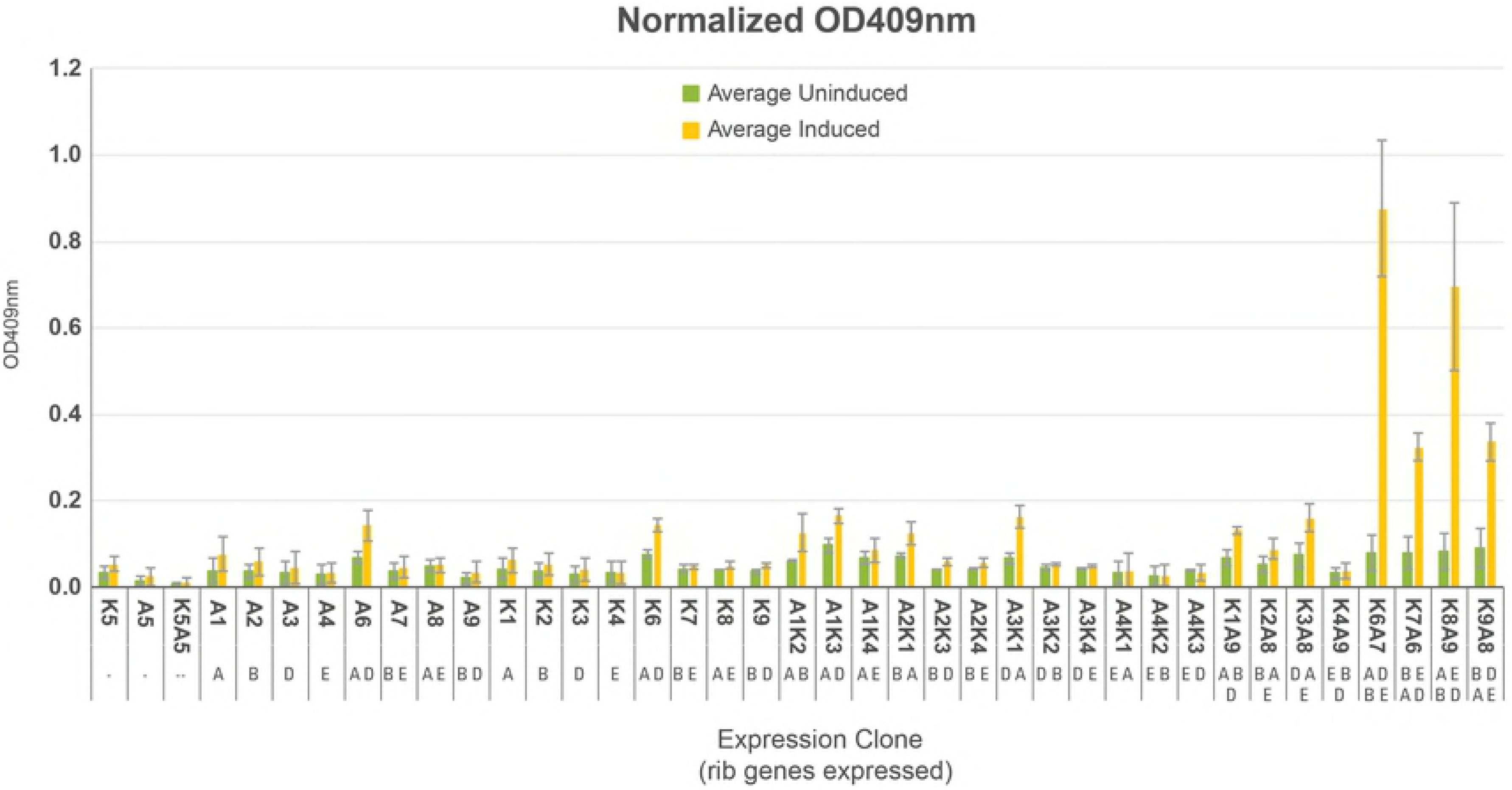
(a) Total Quantites of DMRL Synthesis from *Devices* and *Systems*. Single colonies from *devices* and *systems* listed in Table 3 were purified by re-streaking onto fresh LB-kan-amp plates without IPTG. Three colonies from each plate were cultured in separate tubes containing 3 ml of LB-kan-amp. One hundred microliters of each culture were added to 4.9 ml of LB-kan-amp and incubated until OD600 reached between 0.3 and 0.5. IPTG was added to a final concentration 0.5 mM and incubation continued for three hours after which OD600 values were re-measured. Cultures were centrifuged to remove cells and OD409 values of supernatants obtained. OD409 values were normalized relative to OD600 values measured post IPTG addition and the resulting numbers compared. DMRL was synthesized exclusively by *systems* containing two *devices* each expressing two rib genes –*systems* K6A7, K7A6, K8A9 and K9A8.

**Figure 7.**
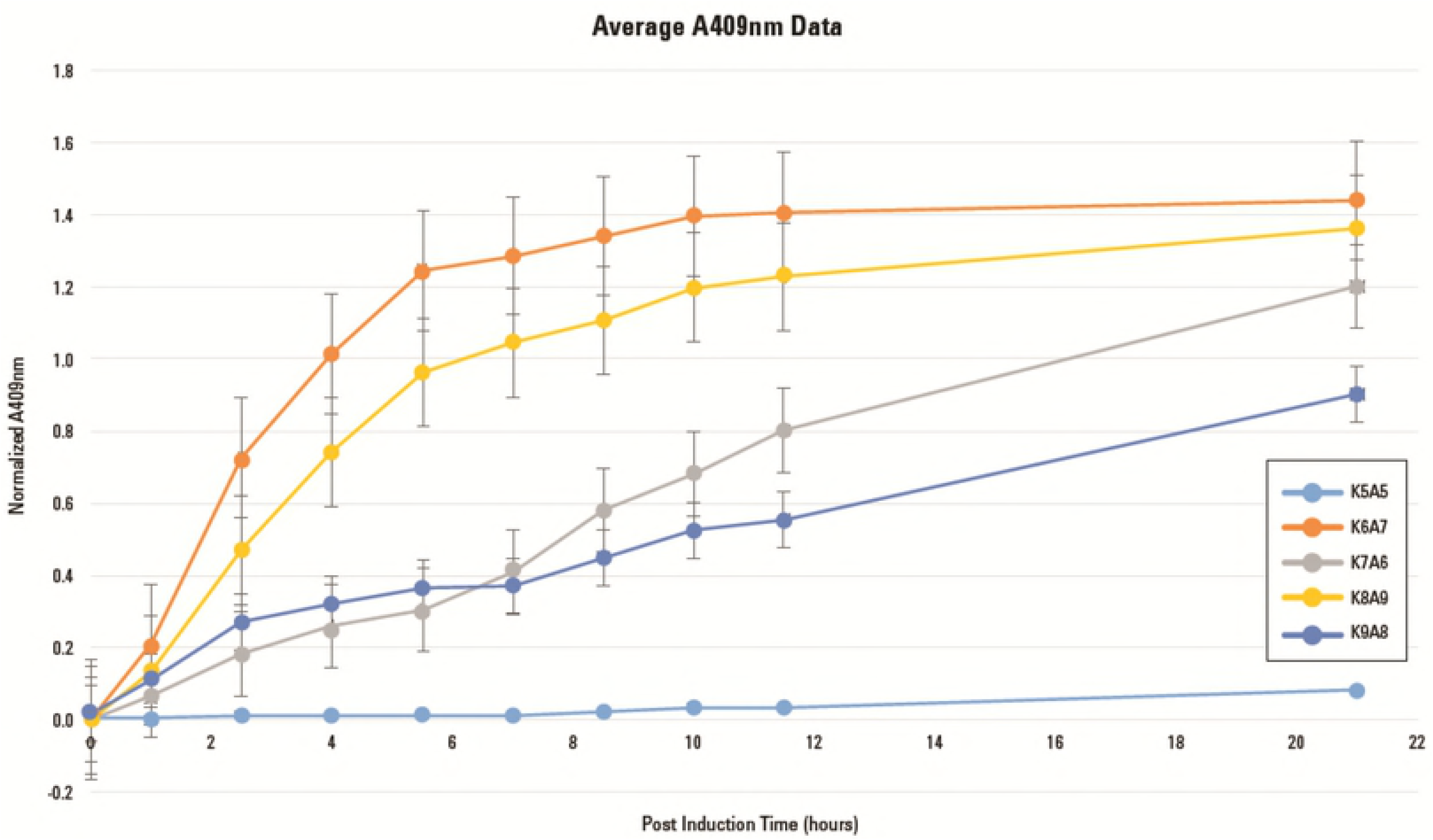
(b) DMRL Synthesis Rates. DMRL synthesis rates of *systems* K6A7, K7A6, K8A9, K9A8 and negative control K5A5 were obtained by inoculating single colonies into 5 ml of LB-kan-amp media followed by overnight incubation at 37°C with shaking at 250 rpm. One ml of these starter cultures was inoculated into separate 250 ml flasks each containing 49 ml of LB-kan-amp. Incubation at 37°C with shaking at 250 rpm continued untilOD600 values reached between 0.3 and 0.5. IPTG was added to a final concentration of 0.5 mM and incubation continued. At regular intervals, 1.0 ml samples were retrieved from each culture and OD600 values measured. Samples were then centrifuged to remove cells and OD409 values of supernatants obtained. OD409 values were normalized by dividing by the OD600 values and these numbers plotted as a function of OD409 post-IPTG addition.

## Discussion

A new method (SureVector) for seamless assembly of biological *<u>parts</u>* into functional *<u>devices</u>* and higher order *<u>systems</u>* is presented that takes advantage of principles learned from prefabrication engineering design and assembly. These are: (1) Functional DNA *parts* are manufactured and quality controlled at a location away from the assembly site (Agilent Technologies, Inc.; (2) *Parts,* other requisite assembly materials and detailed assembly instructions are delivered to the site of construction (the research laboratory) for assembly; (3) from a synthetic biology standpoint, processes (1) and (2) enable rapid and reliable combinatorial assembly of desired *devices* destined for introduction into *E. coli*, mammalian and yeast cells. The salient features of this new method were demonstrated by constructing a set of protein expression *devices* to identify the best *device* for the expression of a target GOI (Nedd5) and a higher order system was constructed to recreate a four-gene biosynthetic pathway *system* (DMRL). The combinatorial assembly power of this process was utilized in developing the DMRL expression *system*. A total of 18 protein expression *devices* were assembled in one day possessing zero, one or two rib gene (bi-cistronic) *parts*. Expression screening was initiated the following day and positive *systems* were selected for structural and functional testing in less than one week. While assembled *devices* were being sequenced for validation of their structural integrity, DMRL was purified and characterized by mass spectrometry. Experiments were also performed to determine optimal DMRL synthesis rates and maximum production. The same type of project out sourced to a third-party vendor would have taken several months to complete (previous experience). While optimizing DMRL production was not the goal of these experiments, it is worth noting that 40 mg of DMRL was purified from 100 ml of media collected after growth of 0.8 g (wet weight) of *E. coli* clone K6A7. In stark contrast, Maley and Plaut (15) required 5 kg of the mold *A. gossypii* to obtain 160 mg of DMRL.

The combinatorial power, simplicity and assembly accuracy of the SureVector process will facilitate building many “multi-device” *systems* including unique biochemical synthetic pathways and novel regulatory circuits. Production of fine chemical intermediates and end-products represents an obvious high-value application.

## Acknowledgement

The authors would like to thank Dr. Bill Webb at The Scripps Research Institute Center for Metabolomics and Mass Spectrometry for his expertise in performing and analyzing all mass spectrometry data.

## Notes

**Funding** This work supported exclusively by Agilent Technologies, Inc.

